# Linear and Nonlinear Oscillatory Functional Brain Networks in Euthymic Bipolar Disorder Classification

**DOI:** 10.64898/2026.01.29.702676

**Authors:** Fatemeh Akrami, Arvin Haghighatfard, Vishal Bharmauria, Tobias Thelen, Amir Hossein Ghaderi

## Abstract

Functional brain network (FBN) dysconnectivity has been repeatedly reported in bipolar disorder (BD). However, it remains unclear how this dysconnectivity manifests from the perspective of oscillatory FBNs, that is, which network measures and frequency bands most reliably capture this alteration. Moreover, it is unknown whether this dysconnection is predominantly expressed through linear or nonlinear interactions.

Here, we investigated properties of oscillatory FBNs in individuals with euthymic BD. Networks were constructed using linear and nonlinear connectivity measures applied to source-localized resting-state electroencephalography (EEG) current density signals. We then quantified whole-FBN and nodal features using conventional and spectral graph theory methods to characterize disorder-related network mechanisms and evaluate their potential as biomarkers.

Significant group differences between BD and control groups were observed in the theta and alpha1 bands. Dynamical whole-FBN alterations were detected primarily in linear oscillatory FBNs, with reduced Shannon entropy and energy in the BD group. These effects were replicated using machine learning, achieving 85% classification accuracy with entropy and energy as the most informative features. In contrast, nodal-level differences emerged mainly in nonlinear FBNs, revealing increased centrality in frontal, and decreased centrality across temporal, and limbic regions.

These findings emphasize distinct, frequency-specific roles of linear and nonlinear oscillatory FBNs in BD, with global dysconnectivity reflected in linear FBNs and local alterations captured by nonlinear connectivity. Moreover, network measures related to synchronization stability and complexity more effectively capture BD-related dysconnectivity, suggesting that dynamic features of oscillatory FBNs may serve as potential biomarkers for BD.

## 1. Introduction

Bipolar disorder (BD) is a chronic psychiatric condition marked by alternating episodes of mania /hypomania and depression, affecting approximately 1–3.9% of the global population[1–4]. Classified under bipolar and related disorders in the Diagnostic and Statistical Manual of Mental Disorders 5 (DSM-5), BD is defined by distinct mood episodes[5]. It is a major contributor to global disability and the overall disease burden[6]. Beyond mood fluctuations, BD is linked to cognitive, emotional, and social impairments, highlighting the need for more effective diagnostic and treatment approaches[7–9].

Pharmacological treatment remains the cornerstone of BD management, with personalized strategies focused on stabilizing acute episodes and preventing relapse[1,10]. Current clinical guidelines emphasize the integration of pharmacotherapy with psychosocial interventions to achieve optimal outcomes[1,11,12]. Psychotherapeutic interventions are generally well-tolerated and associated with minimal side effects[13]. Emerging treatments like transcranial magnetic stimulation (TMS) have shown potential in managing bipolar depression (BD-D), though further research is needed to confirm their efficacy[14]. The limited effectiveness of current therapies underscores the importance of advancing our understanding of brain neurobiology in BD.

From a mechanistic perspective, both functional magnetic resonance imaging (fMRI) and source-localized electroencephalography (EEG)[15–17] studies have revealed that numerous brain regions exhibit aberrant functional activity in individuals with BD compared to healthy control groups. These regions include prefrontal, temporal[15,16], and anterior cingulate cortices, as well as amygdala[18,19], hippocampus[20,21], insula[17], and basal ganglia[22,23]. Importantly, the nature of these functional abnormalities is not static; it demonstrates significant state-dependent alterations, varying according to whether the patient is in a manic, depressive, or euthymic phase[22,24].

Given these extensive body of functional neuroimaging findings implicating diverse brain regions, modules, and connectivity patterns in BD, it is increasingly conceptualized as a disorder of widespread functional brain network (FBN) dysfunction, rather than isolated abnormal activations in specific areas[25,26]. This holistic view posits that BD affects the entire FBN architecture and dynamics[27]. In this context, graph theory analysis of neuroimaging data has emerged as a powerful tool to quantify these global disruptions[28–31]. Studies utilizing this approach have consistently demonstrated altered topological properties in the brain’s connectome of individuals with BD[28,30]. These alterations often include reduced global efficiency[28,31], disrupted integration between specialized FBNs, and a shift towards a less optimal connectivity organization, providing a mathematical framework for the hypothesis that BD is a *dysconnection syndrome* affecting the brain as an integrated system[32,33,33].

However, the established concept of *dysconnectivity* in BD raises several critical questions. The first question that arises is: if BD is a disorder of dysconnectivity, at what level does this dysconnectivity occur? Previous studies have suggested that such dysconnectivity manifests primarily in the linear coupling between activities of different brain regions, that is, in the extent to which simultaneous BOLD signal fluctuations or electrophysiological activities with relatively stable phase relationships reflect coordinated neural communication[25,32,34–36]. However, an open question remains as to whether the brain’s nonlinear dynamic features are also affected by this dysconnectivity. In other words, interactions between regions that exhibit synchronization through more complex, nonlinear phase relationships also become disrupted in BD? This question is of particular importance, as recent studies have demonstrated that nonlinear connectivity measures can be highly informative in characterizing cognitive and affective disturbances in other psychiatric conditions, such as major depression[37], Schizophrenia[38], and attention deficit/hyperactivity disorder (ADHD)[39].

A second pivotal question arises: given the extensive spectrum of neural oscillatory (i.e., delta, theta, alpha, beta) dysfunction in regional activity[40–44] and local functional connectivity[25,41,45–47,47], can global neural dysfunction in BD be considered in all neural oscillatory spectrum as well? Or, at least during euthymic conditions, is it confined to specific oscillatory bands and, consequently, their unique underlying neural systems? Understanding in which oscillatory modes this global functional disconnection occurs can both enhance our comprehension of the neural basis of the BD and the impaired system-level functions, and also contribute to refining quantitative electroencephalography (qEEG) based brain modulation and stimulation protocols, such as EEG neurofeedback[48], transcranial direct current stimulation (tDCS)[49], and TMS[50,51].

Finally, third fundamental inquiry is whether this dysconnectivity manifests as a static topological aberration or as a dynamic impairment in the stability of synchronization between brain regions. If a static, less integrated and probably more segregated topological state is primarily involved, conventional graph theory metrics would be suitable for characterizing this condition[52,53]. Conversely, if the dysfunction is rooted in the instability of synchrony within the FBNs, then a dynamical measure derived from spectral graph theory could serve as a potent biomarker for BD, as it quantifies the temporal variability and oscillatory vigor of the network[37,54–56].

Addressing dynamic characteristics is particularly important, as several recent studies suggest that high-level cognitive-affective features[56–58], as well as network mechanisms, in certain cognitive and affective disorders such as ADHD[59,60] and depression[37,60,61] can be effectively explained by these dynamic stability properties. This goes beyond the neural correlates of these disorders found in the topological features of functional brain networks.

Here, we analyzed EEG signals from individuals with BD in a euthymic phase and from healthy controls using a comprehensive set of methodological approaches. To address the first and second questions jointly, we began by applying source-localized EEG analysis to obtain spatially precise estimates of cortical activity, leveraging recent methodological advances that improve the localization of electrophysiological signals[37,62,63]. Using these source-level signals, we constructed FBNs with both linear (lagged coherence) and nonlinear (phase synchronization) connectivity metrics to determine which type of coupling more accurately characterizes dysconnectivity in BD. We then quantified whole-FBN metrics and regional centrality features within linear/nonlinear FBNs to determine which connectivity framework (linear or nonlinear) more clearly differentiated individuals with BD from healthy controls across both large-scale organization and hub-level contributions.

To address the third question and to determine whether topological properties or dynamic synchronization features of a network better reflect neural dysconnectivity in BD, we used several network science methods: 1-Conventional graph theoretical analyses by applying clustering coefficient (CC) and global efficiency (Ef) to assess segregation and integration, respectively[29,64,65]. 2-Spectral graph theory by using graph energy (H) to evaluate synchronization stability[54,56], and 3-information-theoretic analysis by measuring Shannon entropy (S) of graph to examine the complexity of connectivity patterns[54,56].

We expected that the application of nonlinear connectivity measures would reveal distinct patterns compared to those obtained from linear connectivity analyses. We hypothesized that FBN measures would capture dysconnectivity across multiple frequency bands, with these alterations reflected in both topological and dynamical properties of the networks. Given the inherent complexity of neural dynamics in BD, we further anticipated that features representing FBN dynamics and complexity (e.g., H and S) would serve as stronger predictors for distinguishing BD participants. Through machine learning (ML) analyses, these FBN-derived features could provide novel EEG-based biomarkers of BD, grounded in source-localized functional connectivity.

## 2. Materials and Methods

### 2.1. Participants

Forty-seven individuals participated in this study, including 23 participants diagnosed with BD (mean age = 30.81 years; 10 males, 14 females) and 24 healthy controls with no history of psychiatric or neurological disorders (mean age = 29.47 years; 11 males, 14 females). BD diagnoses were confirmed using the Structured Clinical Interview for DSM-5 (SCID-I) by a board-certified psychiatrist [43,66].

At the time of assessment, all BD participants were in euthymic phase. All participants were screened to exclude histories of serious head injury, neurological or psychiatric conditions (other than BD in the patient group), and any substance or alcohol use within two weeks before EEG recording. Vision and hearing were confirmed to be normal or corrected-to-normal for all participants.

The study was approved by the ethics committee of Islamic Azad University, Tehran, Iran (IR-1398.093) and conducted in accordance with the declaration of Helsinki. All participants provided written informed consent following a detailed explanation of the study procedures.

Following EEG data acquisition and preprocessing, seven participants (three BD, four controls) were excluded due to excessive EEG artifacts. The final sample comprised twenty participants in each group.

### 2.2 EEG Acquisition and Preprocessing of Signals

Resting-state EEG data were recorded for 5 minutes in the eyes open condition using a Brainmaster Discovery-24 amplifier. EEG signals were collected from 21 electrodes at a sampling rate of 250 Hz with 24-bit resolution. Nineteen scalp electrodes (Fp1, Fp2, F3, F4, C3, C4, P3, P4, O1, O2, F7, F8, T3, T4, T5, T6, Fz, Cz, Oz) were positioned according to the international 10–20 system. Two additional electrodes (A1 and A2) placed on the preauricular areas served as a linked-ear reference. Participants were instructed to keep their eyes open, minimize blinking, and fixate on a target to maintain a stable resting state.

The offline preprocessing of EEG data was conducted using the EEGLAB toolbox[67] in MATLAB R2024b. EEG signals were bandpass filtered between 0.1–40 Hz using a fifth-order digital filter. Artifact rejection involved both automatic and manual approaches. The continuous data were segmented into 4-second epochs. Specifically, automatic detection was carried out using the channel statistics tool in EEGLAB in combination with the artifact subspace reconstruction (ASR) algorithm. ASR identifies clean calibration data segments and estimates the standard deviation of PCA-extracted components, while excluding physiological activity such as alpha and theta rhythms through filtering. Data segments are then rejected if they exceed a threshold defined relative to the standard deviation of the calibration data (default setting)[68]. This approach helps to exclude EEG segments containing physiological artifacts including eye movement/blinks, body movement or muscle activity. Finally, all data underwent manual visual inspection to ensure exclusion of residual noisy segments and to confirm the quality of the retained epochs. Cleaned EEG data were exported for further source localization and network analyses.

### 2.3. Standard LORETA source localization algorithm and functional connectivity analysis

Standardized low resolution brain electromagnetic tomography (sLORETA) algorithm[69] is an inverse solution method used to estimate the cortical sources of scalp-recorded EEG activity by solving the inverse problem under the assumption that neighboring neuronal sources have highly similar activity[70]. It provides a smooth spatial distribution of the estimated sources and estimates the distribution of current density throughout the cortex without requiring a priori assumptions about the number or location of sources. The sLORETA algorithm has been validated for accurate source estimation in different EEG studies[71–74]. Moreover, previous studies have shown that 19-channel EEG can provide accurate estimates of EEG source current densities [57,75,76].

Using sLORETA, we computed current density values across all artifact-free epochs (extracted from EEGLAB) in 84 regions of interest (ROIs), representing the maximum number of available regions in the LORETA-Key software. These ROIs include 42 cortical Brodmann areas (BAs) in the left hemisphere and 42 cortical BAs in the right hemisphere, covering BAs 1–11, 13, 17–25, and 27–47.

### 2-4 Functional connectivity and Adjacency matrices

Functional connectivity was assessed between all pairs of ROIs (84 cortical BAs). Connectivity analyses were conducted separately for seven frequency bands: delta (1–4 Hz), theta (4.5–7.5 Hz), alpha1 (8–10 Hz), alpha2 (10.5–12 Hz), beta1 (12.5–18 Hz), beta2 (18.5–21 Hz), and beta3 (21.5–30 Hz). We calculated two different functional connectivity measures, lagged coherence and nonlinear connectivity, which evaluates linear and nonlinear coactivations between brain regions respectively. These measures were calculated using the sLORETA connectivity toolbox [64].

Lagged coherence estimates the functional coupling between two brain regions by separating the imaginary part of the cross-spectrum, thereby minimizing volume conduction effects[77]. The lagged coherence between two time series, *x* and *y*, at a frequency ω is given by:

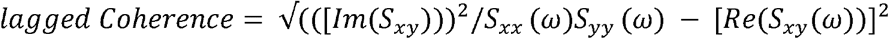

where *ω* represents the frequency, *S*_*xy*_(*ω*) denotes the cross-spectral density between time series *x* and *y*, and *S*_*xx*_(*ω*)and *S*_*yy*_(*ω*) correspond to the auto-spectral densities of each time series. The terms *Im* and *Re* refer to the imaginary and real components of the cross-spectrum, respectively.

To detect higher-order dependencies and dynamic interactions between regions not captured by linear metrics (e.g., lagged coherence), we computed nonlinear connectivity using phase-based measures[78]. These quantify the consistency of instantaneous phase relationships between signals, capturing synchronization beyond simple linear correlations. In this study, nonlinear connectivity was quantified using sLORETA’s Lagrangian connectivity (LC) measure which is defined as[79]:

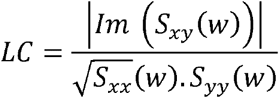

where *S*_*xy*_ is the cross-spectral matrix of sLORETA-estimated cortical current densities. This metric isolates nonlinear phase synchronization by first utilizing the imaginary component of the cross-spectrum to reject volume conduction artifacts, then operating on source-level time series derived from the weighted minimum norm solution and finally detecting the time-lagged interactions that reflect genuine neuronal coupling.

We then used these undirected connectivity measures to construct separate linear and nonlinear undirected weighted FBNs. In this approach, each brain region was considered as a *node* and the functional connectivity values between regions were represented as *edges*[52,80]. For each participant in each group, two FBNs (linear and non-linear) were constructed as separate undirected weighted adjacency matrices (in MATLAB R2024a). To ensure that the network measures were not driven by the mean connectivity values and truly reflected the dynamics, complexity, and topology of the networks, we constructed null models for each adjacency matrix. This model generated randomized networks by shuffling the upper triangular elements of the original matrix while preserving symmetry and avoiding self-connections. These null models served as a baseline for comparison. The resulting adjacency matrices, alongside their null-model counterparts, were subsequently used for graph theoretical analysis to investigate dynamical and topological brain network features[81,82].

### 2.5. Network analysis

#### 2.5.1 Whole functional brain network measures: topological and dynamical

Following the construction of adjacency matrices, we calculated topological, and dynamical measures of functional brain networks. We first average over all connection weights in each adjacency matrix and calculated the connection density (CD) for each participant. Then, we applied approaches from conventional graph theory analysis to evaluate key topological features, CC, and Ef, which reveal the network segregation and integration, respectively.

CC measures the extent of local connectivity within the network, reflecting the degree of functional segregation. Networks with highly interconnected nodes demonstrate higher CC values, indicating increased local information processing within the brain[37,83]. Mathematically, for a weighed undirected network, the CC of node *i* is defined as[84]:

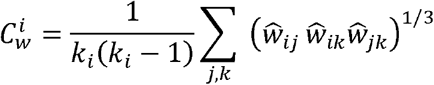

where, *K*_*i*_ is the degree of node *i, ŵ*_*ij*_ *ŵ*_*ik*_ *ŵ*_*jk*_ are edge weights normalized by the network’s maximum weight 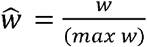. The ∑_*j,k*_ runs over all pairs of neighbors connected in a triangle (*i* − *j* − *k* − *i*)The exponent 1/3 computes the geometric mean of triplet weights. We used *clustering_coef_bd* function from brain connectivity toolbox [80] in MATLAB R2024a to compute this measure.

Ef quantifies the network’s capacity for information integration across distant brain regions, representing the efficiency of communication within the network. Networks with high Ef values facilitate faster information transfer, indicating higher functional integration [37]. Mathematically, for a weighed undirected network, *Ef* is defined by [80]:

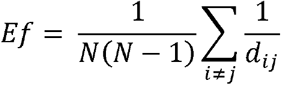

where *d*_*ij*_ is the shortest path length between nodes *i* and *j* which is derived from inverse edge weights. We used efficiency_wei function from brain connectivity toolbox [80] in MATLAB R2024a to compute this measure.

In addition to conventional graph theory measures, we applied spectral graph and information theory analyses to investigate network wide dynamic properties and complexity of connectivity patterns. Specifically, we computed H and S to assess satiability of synchronization and network complexity, respectively.

H is related to eigenvalue spectrum of adjacency matrix and represents the stability of functional synchronization between brain regions[54,74]. Mathematically, for a weighed undirected adjacency matrix, H is defined by the sum of absolute eigenvalues[54]:

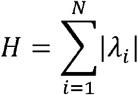

where *λ*_*i*_ are eigenvalues of a given adjacency matrix. We used a self-developed function in MATLAB R2024a to calculate H.

S is inversely related to the probability of state occurrence, where minimal entropy arises when all connections have uniform strength (i.e., higher probability) and maximal entropy occurs when the network exhibits diverse, less predictable connectivity patterns. Increased entropy is typically associated with more complex and dynamic neural interactions[54]. Mathematically, for a weighted undirected adjacency matrix W, S is defined by[54]:

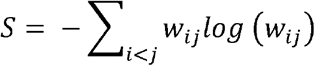

Where weights are normalized to sum to unity. We used an *entropy* function in MATLAB R2024a to calculate S.

#### 2.5.2. Centrality Measures

Centrality measures are essential for identifying the most influential nodes within FBNs, as they reflect the role of specific brain regions in facilitating communication and information flow. These measures are particularly valuable in understanding network alterations in psychiatric conditions, where localized disruptions in connectivity may influence overall brain network organization. Here, to evaluate both local and global regional involvement of each node in the FBN, we used two centrality measures i.e., eigenvector centrality and betweenness centrality, which assess local and global hubness respectively[37,80].

*Betweenness centrality* (BC) quantifies the number of shortest paths which are crossing a node and can assess the influence of a node in controlling the global information flow within the FBN. Nodes with high BC lie on numerous shortest paths between other nodes, giving them a critical role in wide-range inter-regional communication. This measure highlights regions that may act as global hubs for information flow in the FBN. Higher BC suggests that a node has substantial control over the network’s information transfer, while lower values indicate more peripheral roles [80]. In this study, BC was calculated for all the nodes. To calculate BC, we used efficiency_wei function from brain connectivity toolbox [80].

Eigenvector centrality (EigC) is related to eigenvector values (corresponding to largest eigenvalue) of adjacency matrices and quantifies the influence of a node within a network by considering both its direct connections and the significance of its neighboring nodes. Nodes with higher EigC values are more strongly connected to other surrounding influential nodes, highlighting their role in facilitating information flow across in the local scale[85]. Unlike BC, which primarily captures pathways of information transfer, EigC emphasizes a node’s influence within the communities or local clusters[86]. We calculated EigC for each node using *eigenvector_centrality_und* function from the brain connectivity toolbox [80].

#### 2-5-3 Null models for graph theoretical measures

All graph theory analyses for calculating CC, CPL, H, BC, and EigC were also performed on the randomly shuffled matrices (null models) for each participant[87]. The values obtained from the original matrices were then divided by (for CC, CPL, and H) or subtracted from (for BC and EigC) the values derived from the null matrices to ensure that these measures were dependent on network topology or dynamics and not influenced by potential differences in connection density. Since entropy values depend on the distribution of connection weights and remain unchanged by their spatial rearrangement, this measure was not normalized with null models.

### 2.6. Statistical Analysis

To find the statistically significant differences between the BD group and control group in FBN properties, and regarding the limited number of our sample size, we employed nonparametric permutation tests. This non-parametric approach will help to reduce the statistical errors which may rise using parametric statistical analysis, particularly when the assumptions of normality and homogeneity of variance are not met in small sample sizes[88].

The number of random shuffles required for a permutation test depends on the specific test and the desired significance level. In our analysis, we determined that 5000 random shuffles were necessary to achieve a normal distribution in t-values with a significance level of p = 0.05. To control for type I errors when comparing five whole-FBN measures across seven frequency bands (35 comparisons in total), we applied a false discovery rate (FDR) correction[89,90].

To compare the regional EigC and BC (in all 84 BAs using nonparametric permutation tests in the frequency bands where significant differences in the whole-FBN measures were observed (two frequency bands). These multiple comparisons were also corrected using FDR. The whole statistical analyses were separately performed for two sets of linear and non-linear matrices.

### 2.7 Machine Learning Analysis

To investigate whether whole-FBN features could effectively distinguish between the BD and control groups, and to evaluate potential network-based biomarkers for BD, we applied machine learning (ML) classification models. Specifically, we used a supervised support vector machine (SVM) due to its effectiveness in handling complex, high-dimensional data and the possibility for its suitability for small sample sizes[91].

SVM identifies the optimal hyperplane to maximize the separation between data points of different classes. By maximizing the margin between support vectors (the most influential data points) and the decision boundary, SVM minimizes classification errors. We employed MATLAB’s *fitckernel* function to implement a kernel-based SVM. The Kernel scale was automatically determined by the function to define the influence of each data point in the transformed higher-dimensional space. Regularization value (Lambda) was initially auto determined (≈0.025) and further adjusted to 0.01, 0.05, and 0.1.

The SVM model was trained using five whole-FBN measures (CD, CC, CPL, H, and S) as predictive features for classification. To enhance generalizability and minimize the risk of overfitting, we applied 5-fold cross-validation, dividing the dataset into five equal folds, training the model on four folds, and validating it on the remaining fold[92,93]. Due to the limited sample size, we further improved reliability by repeating the cross-validation process five times with different initial random shuffles of the data.

To evaluate classification performance, confusion matrices were presented to show true positives, true negatives, false positives, and false negatives. We also calculated the receiver operating characteristic (ROC) curve and the area under the curve (AUC) for each fold to assess model discrimination. To provide a more detailed assessment of model performance, we also used precision-recall area under the curve (PR-AUC) which considers the potential class imbalance. The average accuracy across all folds was also computed. To identify the most discriminative features for classification, we performed feature selection using the ReliefF algorithm [94] implemented in MATLAB R2024a. This approach ranks features based on their ability to distinguish between classes while accounting for feature dependencies. The entire methodological pipeline is presented in Figure 1.

**Figure 1.**
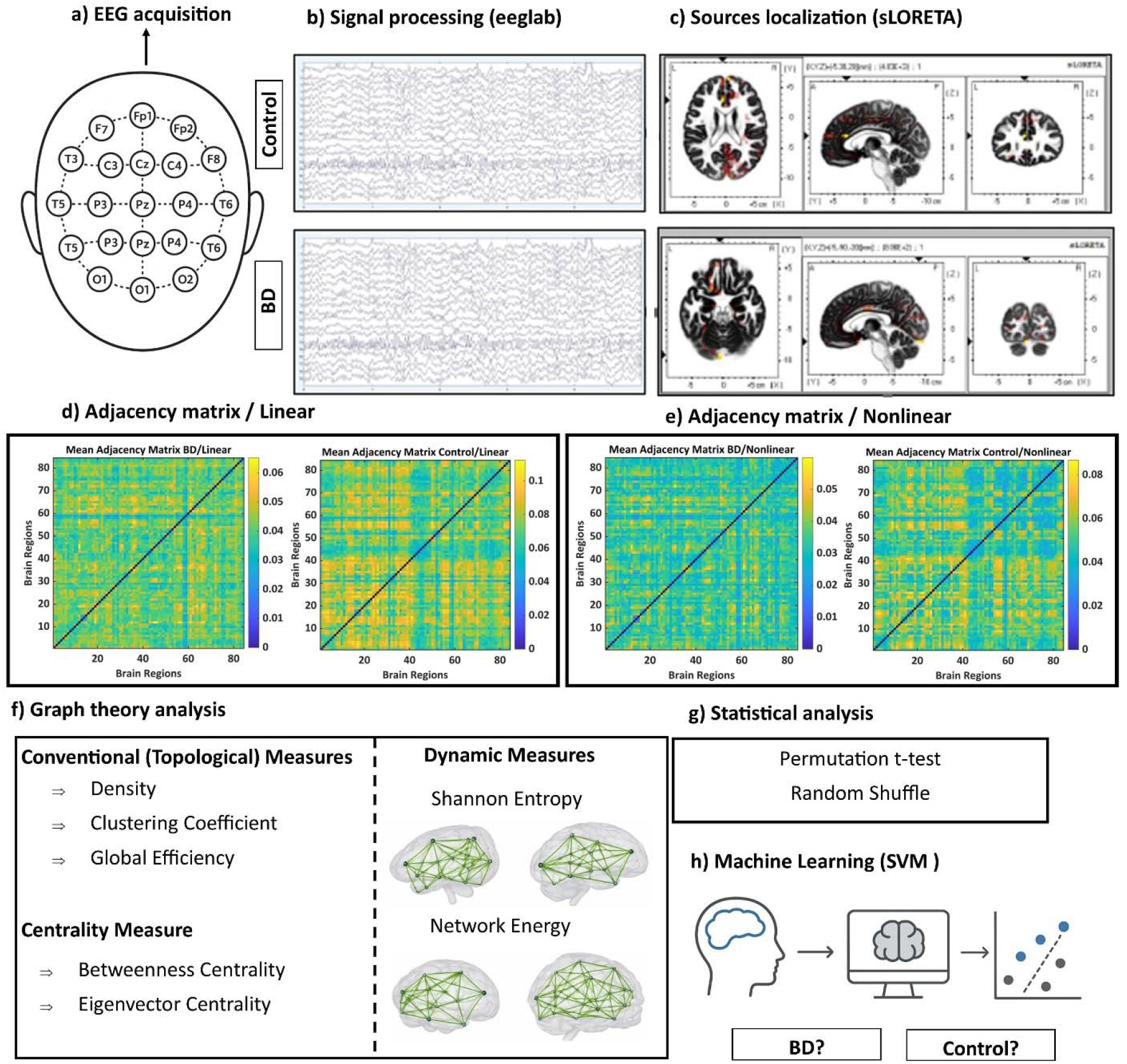
Methodological pipeline of the study. The analysis consisted of eight sequential steps: (a) EEG signals were acquired using the standard 10–20 electrode placement system. (b) signal preprocessing and artifact removal were performed using EEGLAB; (c) cortical source activity was reconstructed using sLORETA. (d) Linear functional connectivity matrices were constructed to represent inter-regional brain interactions. (e) Nonlinear functional connectivity matrices were constructed for both groups to capture complex inter-regional interactions. (f) Graph-theoretical analysis was performed, including topological measures (connection density, clustering coefficient, and global efficiency), nodal centrality measures (betweenness and eigenvector centrality), and dynamic measures (Shannon entropy and network energy). (g) Statistical analysis was conducted using permutation t-tests with random shuffling. (h) Machine-learning classification was carried out using a SVM trained on the extracted features to discriminate between BD and control groups.

## 2. Results

We first compare the whole-FBN features between the BD and control groups using permutation t-test followed by FDR correction. As shown in Figure 2, across whole-FBN measures, significant differences between BD and control groups were observed primarily in lower frequency bands (theta and alpha1) and mostly in networks constructed using linear connectivity measures. Specifically, with linear connectivity, significant differences between BD and control groups were observed in alpha1 CC (t = 3.54, p = 0.001), theta H (t = 2.8, p = 0.03), and alpha1 S (t = 4.1, p = 0.001). With nonlinear connectivity, the only significant difference between groups was observed in alpha1 S (t = 2.8, p = 0.03), which was also significant with linear connectivity, see Figure 3.

**Figure 2.**
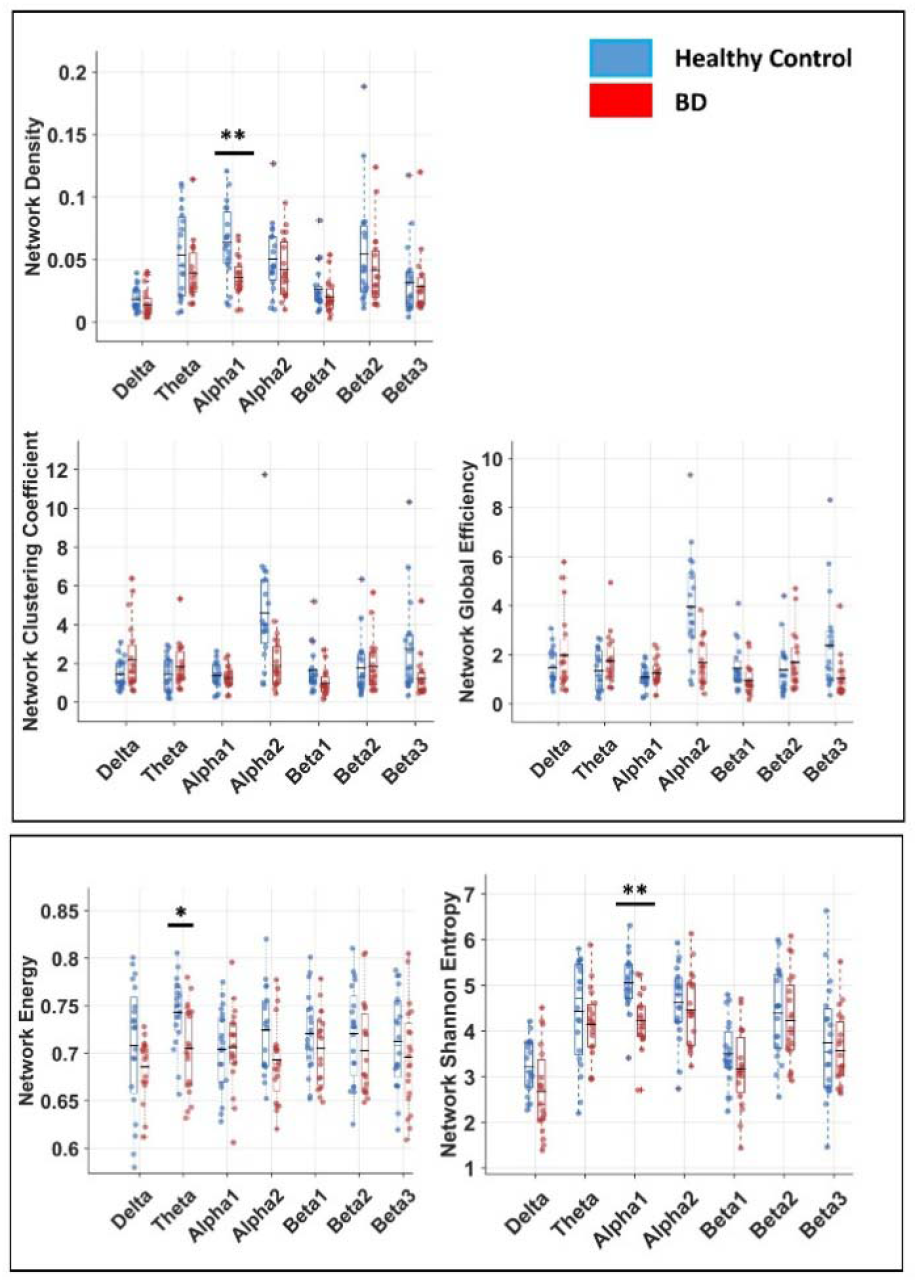
Linear functional connectivity network measures across frequency bands.Boxplots illustrating graph-theoretical network measures derived from linear lagged coherence across seven frequency bands (Delta, Theta, Alpha1, Alpha2, Beta1, Beta2, Beta3) for healthy controls (blue) and BD (red) groups. The panels display (from top to bottom) network density, CC, GE, H, and S. Each dot represents an individual participant. Statistically significant group differences identified using permutation t-tests with FDR correction are indicated by asterisks (*p < 0.05; **p< 0.01).

**Figure 3.**
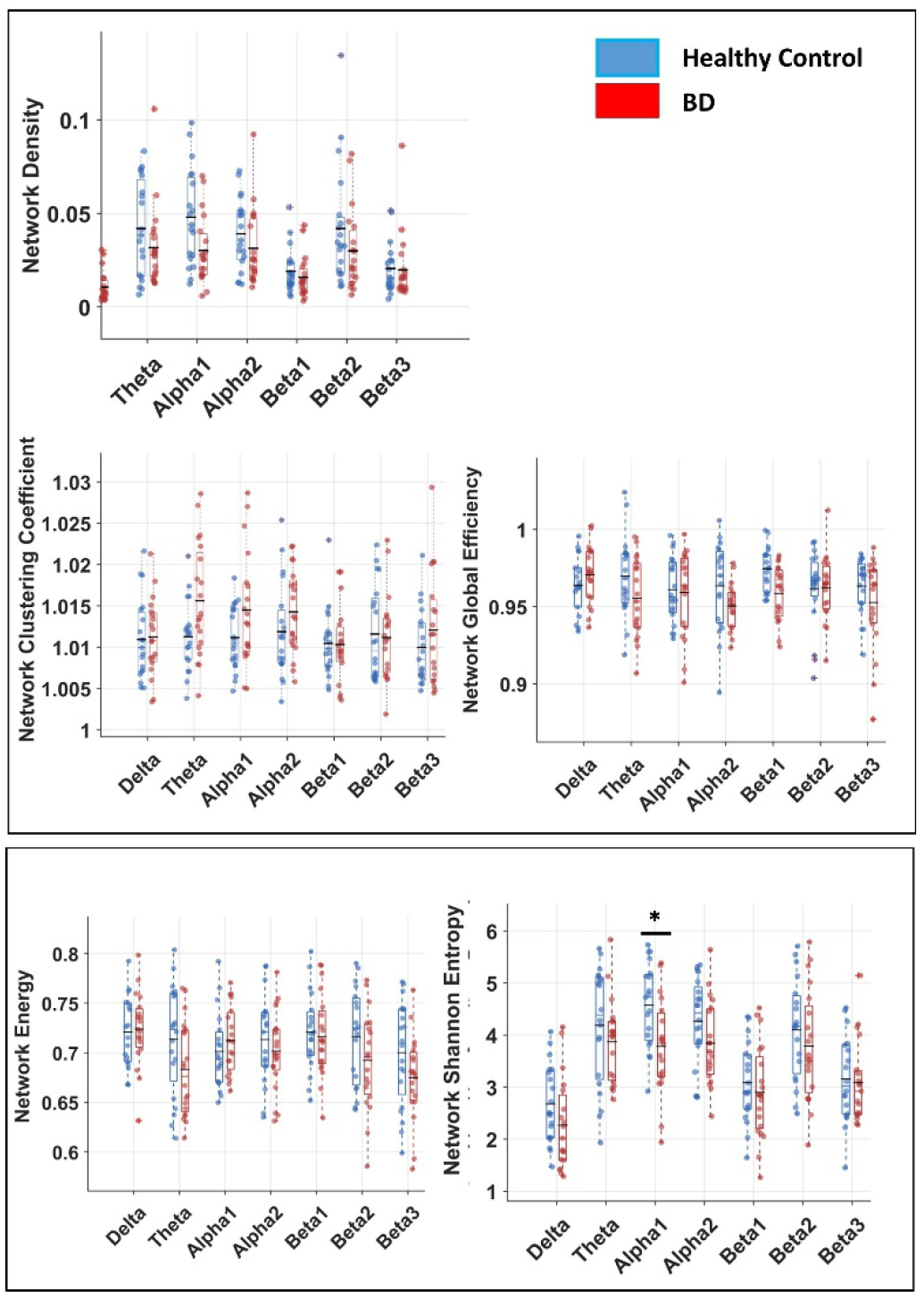
Nonlinear functional connectivity network measures across frequency bands. Boxplots illustrating graph-theoretical network measures derived from nonlinear lagged coherence across seven frequency bands (Delta, Theta, Alpha1, Alpha2, Beta1, Beta2, Beta3) for healthy controls (blue) and BD (red) groups. The panels display (from top to bottom) CD, CC, GE, H, and S. Each dot represents an individual participant. Statistically significant group differences identified using permutation t-tests with FDR correction are indicated by asterisks (*p < 0.05).

Subsequently, we examined centrality metrics to determine the most influential nodes (brain regions) within the networks across frequency bands. After FDR correction, significant group differences in EigC were found in the theta band under the nonlinear connectivity condition. Specifically, several right-hemispheric regions showed altered EigC values (Figure 4). In comparison with the control group, the BD group exhibited significantly higher EigC in BA 8 (frontal eye field; t = 3.14, p = 0.04), while they showed lower EigC values in the following regions: BA 20 (inferior temporal gyrus; t = -3.75, p = 0.04), BA 21 (middle temporal gyrus; t = -3.18, p = 0.04), BA 22 (Wernicke’s area; t = -3.53, p = 0.04), BA 28 (ventral entorhinal cortex; t = -3.18, p = 0.04), and BA 36 (perirhinal cortex; t = -3.13, p = 0.04).

**Figure 4.**
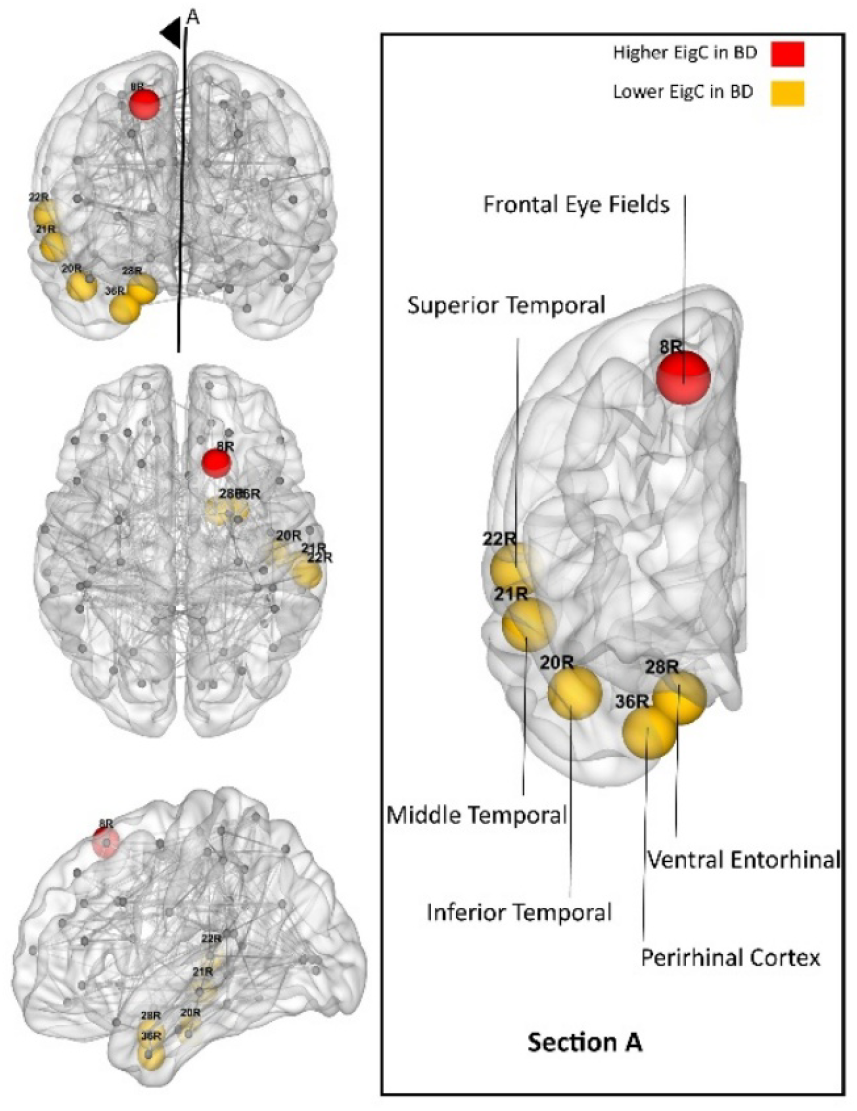
Brain regions showing significant group differences in EigC in the Theta band under the nonlinear lagged coherence condition. Colored nodes indicate regions with altered EigC in the BD group compared to controls. Red indicates higher EigC in the BD group, and yellow indicates lower EigC.

Thereafter, to evaluate whether the whole-FBN measures could serve as biomarkers for discriminating the BD group, we applied ML approaches. Specifically, separate SVM models were trained using four whole-FBN measures: CD, CC, Ef, H, and S in the different frequency bands.

In the Alpha1 frequency band, the SVM classifier demonstrated strong performance in distinguishing between BD and control groups. Model performance was assessed using a 5-fold cross-validation approach. In general, the best performance was achieved an accuracy of 85%, correctly classifying all 20 control participants and 14 out of 20 BD participants (Figure5.a). The hyperparameters for this accuracy were lambda = 0.025, kernel scale = 1. The AUC for both control and BD groups was 79%, indicating a strong ability of the model to distinguish between the two groups. Precision-Recall AUC (PR-AUC) values provided a more nuanced view: 75% for the control group and 87% for the BD group (Fig 5.b). The higher PR-AUC for the BD group suggests the enhanced capability of the model in identifying individuals with BD.

**Figure 5.**
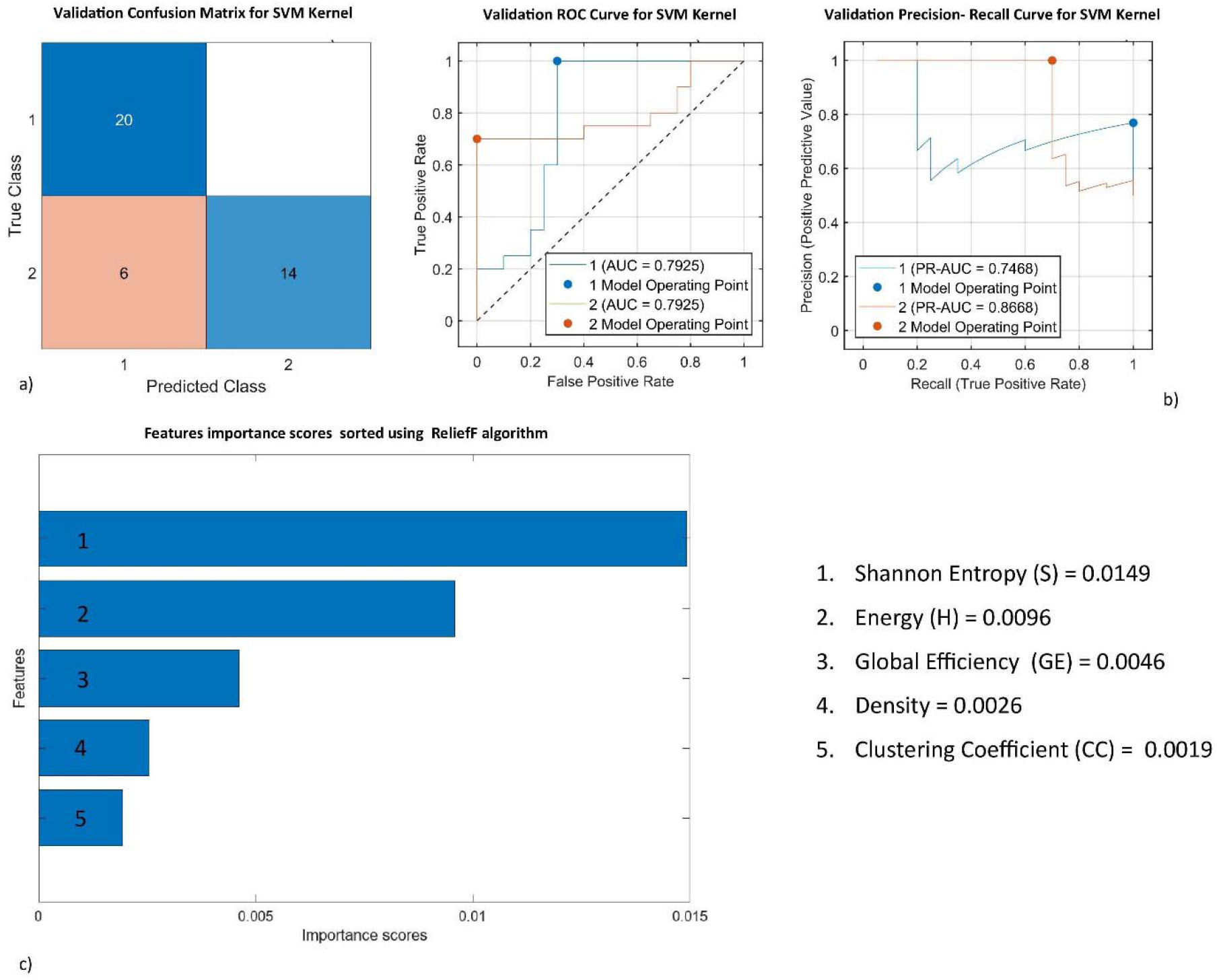
Performance evaluation and feature importance for the SVM kernel model used to classify BD and control groups based on graph-theoretical network measures. (a) Confusion Matrix: The model correctly classified 20 control and 14 BD participants, with 6 misclassified BD cases. (b) ROC and Precision–Recall Curves: The area under the ROC curve (AUC) was 0.7925 for both classes. The Precision–Recall AUC (PR-AUC) was 0.7468 for the control group and 0.8668 for the BD group. (c) Feature Importance: Ranked using the ReliefF algorithm, with Shannon Entropy (S) having the highest importance (0.0149), followed by Energy (H), Global Efficiency (GE), Density, and Clustering Coefficient.

To identify the most discriminative features, we computed feature importance scores using a ReliefF algorithm[95,96] for feature selection. As shown in Figure 5.c, S showed the highest discriminative power, followed by H. In contrast, Ef, CD, and CC exhibited relatively lower importance scores. These findings suggest that features related to FBN dynamics and complexity (S and H) are more critical for classification in this context than topological network metrics.

## Discussion

### Summary of hypotheses and main findings

This study aimed to characterize FBN dysconnectivity in BD by examining whether network alterations extend beyond linear interactions to nonlinear dynamics, whether they are frequency-specific, and whether they primarily reflect static topological changes or dynamic instability of synchronization. Significant group differences were mainly observed in the theta and alpha1 bands. At the whole-FBN level, individuals with BD showed reduced S and CD in the alpha1 band, along with reduced graph H in the theta band, indicating lower global connectivity strength, reduced complexity, and decreased stability of functional synchronization. In contrast, classical topological measures showed limited sensitivity to group differences. At the regional level, nonlinear theta-band networks revealed enhanced hub centrality in the right dorsolateral prefrontal cortex (BA 8) and reduced centrality in temporal and limbic regions, suggesting an altered frontal–temporal–limbic balance in BD. Importantly, ML analyses demonstrated that dynamic and complexity-related network features, particularly S and H, provided the highest discriminative power, supporting their potential as EEG-based biomarkers for BD.

### Superiority of linear connectivity measures and the complementary role of nonlinear metrics

The present findings indicate that linear connectivity measures, particularly coherence-based metrics, are more sensitive to whole-FBN alterations in BD than nonlinear alternatives. Linear measures such as correlation and coherence have been widely used to construct FBN and have consistently revealed disrupted connectivity patterns in BD across EEG and fMRI modalities [22,25,97–102]. In line with this literature, group differences in the present study were detected more robustly in networks derived from linear connectivity, suggesting that dysconnectivity in euthymic BD is predominantly expressed through altered linear coupling between distributed neural populations.

This pattern closely aligns with the coherence hypothesis, which posits that effective neural communication relies on in-phase, stable, and non-random phase relationships across oscillatory activity in distributed brain regions[103–106]. Linear coherence is particularly well suited to capture such frequency-specific synchronization and has been shown to underlie a broad range of cognitive and affective functions, including attention, perception, working memory, and cognitive control[104,107–111]. Importantly, our results reinforce the relevance of linear synchronization as a core mechanism of network dysfunction in BD which is consistent with previous studies[22,27,66,99].

While linear connectivity measures demonstrated greater sensitivity to group-level differences in whole-FBN features, this does not diminish the relevance of nonlinear connectivity metrics. Rather, our findings suggest that nonlinear measures capture complementary aspects of brain network organization that are not fully accessible through linear synchronization alone. Nonlinear connectivity is specifically designed to detect higher-order and more complex dependencies between neural signals, including interactions that are not characterized by stable and non-random phase relationships, and may reflect transient, state-dependent, or cross-scale coordination mechanisms within functional brain networks[112,113].

In the current results, nonlinear connectivity measures were particularly informative at the regional (hub-level) scale, where alterations in EigC were detected in frontal, temporal, and limbic regions. This pattern indicates that nonlinear metrics may be more sensitive to local reweighting of hub influence and hierarchical organization, even when whole-FBN alterations are dominated by linear coupling. Such regional specificity is consistent with the notion that nonlinear interactions support flexible, context-dependent communication, which may remain partially preserved or compensatory in euthymic bipolar disorder.

Taken together, these observations support a multilayer framework for FBN analysis, in which linear connectivity captures dominant, frequency-specific mechanisms of stable synchronization underlying whole-FBN integration, whereas nonlinear measures provide additional insight into fine-grained, adaptive, and hierarchical network dynamics. From this perspective, nonlinear connectivity does not replace linear coherence but extends it, enabling a more complete characterization of network dysfunction in BD.

### Frequency-specific oscillatory network alterations in bipolar disorder

A major strength of the present study lies in its ability to investigate oscillatory FBNs by combining source-resolved EEG with graph-theoretical analyses. This approach uniquely bridges two traditionally separate perspectives in network neuroscience: the spatially resolved network framework commonly employed in fMRI studies and the direct measurement of neuronal electrophysiological activity provided by EEG [37,57,75,114]. By reconstructing cortical source activity and analyzing frequency-specific interactions between brain regions, the current framework enables the examination of FBNs with anatomically defined nodes while preserving the millisecond-scale temporal resolution necessary to capture neural oscillations and their synchronization dynamics.

Our results suggested significant group differences were predominantly confined to the theta and alpha1 bands, indicating that global dysconnectivity in euthymic BD does not uniformly affect the entire oscillatory spectrum but instead targets specific neural rhythms. This observation is consistent with a growing body of EEG literature suggesting that large-scale network dysfunction in BD is selectively expressed within specific oscillatory systems, rather than reflecting a broadband disruption of neural activity[50,115,116].

Alterations in the theta band were primarily reflected in reduced stability of network synchronization and changes in regional hub centrality, particularly within frontal, temporal, and limbic areas. Theta oscillations are widely implicated in long-range communication, affective regulation, and coordination between prefrontal and limbic systems[117], processes that are consistently disrupted in BD[118–121]. Consistently, previous EEG studies have reported abnormal theta power and connectivity in BD[46,122–124].

In the alpha1 band, reductions in global connectivity and network complexity were observed across both linear and nonlinear connectivity frameworks. Alpha oscillations, particularly in the lower alpha range, are thought to play a key role in large-scale functional integration[125], inhibitory control[126], and the regulation of information flow across cortical systems[127]. A substantial body of previous EEG research has demonstrated that BD is associated with marked abnormalities in alpha-band activity, even during euthymic states [128–131]. These studies have consistently reported alterations in alpha power, alpha reactivity, and alpha coherence, suggesting a fundamental disruption in the oscillatory mechanisms that support resting-state brain organization in BD [128–131].

### Differential discriminative power of graph-theoretical measures between groups

Beyond demonstrating the presence of oscillatory FBN alterations in BD, an important contribution of the present study lies in clarifying which classes of graph-theoretical measures are most sensitive to group differences and what these differences imply about the underlying neurobiological mechanisms. Our results show that measures capturing network dynamics and complexity, rather than classical topological indices, provided the strongest discrimination between individuals with BD and healthy controls.

Conventional graph-theoretical metrics such as CC and Ef are designed to characterize relatively static properties of network architecture, including segregation and integration[52,54,80]. Although these measures have been widely applied in BD research and have revealed dysconnectivity, including altered small-world organization and reduced efficiency in several fMRI and EEG studies[26,32,66], their sensitivity in the present dataset was limited. Importantly, we observed a significant reduction in CD in the BD group, indicating a lower overall level of functional coupling across the network. At a basic level, this finding directly supports the dysconnectivity hypothesis in BD which suggests BD is associated with a global reduction in functional interactions among distributed brain regions.

However, many previous graph-theoretical studies have not explicitly accounted for differences in CD by employing appropriate null models. This is a critical methodological consideration, as reduced density alone can trivially lead to lower values of other topological measures, such as CC and Ef, without necessarily reflecting genuine alterations in network organization[87]. In the present study, we therefore applied density-controlled null models to disentangle true topological effects from those driven by overall connectivity strength. After this normalization, CC and Ef no longer showed reliable group differences. This pattern suggests that previously reported alterations in these metrics may, at least in part, reflect underlying differences in network density rather than fundamental changes in the brain’s static topological architecture. Consequently, our findings indicate that during the euthymic phase, BD may not primarily involve large-scale reconfiguration of network topology, but rather a reduction in overall coupling combined with impairments in the dynamic organization and stabilization of functional interactions.

In contrast, S and H emerged as the most discriminative whole-FBN features, both at the statistical level and in the machine learning framework. S, which reflects the diversity and unpredictability of connection weight distributions, was significantly reduced in the BD group, particularly in the alpha1 band, and was identified as the most informative feature in the SVM classifier.

Reduced S can be interpreted as a marker of decreased functional complexity and a reduced repertoire of network states, indicating that the brain operates within a narrower dynamical range[54]. In the context of BD, this reduction in complexity further suggests that global network connectivity becomes less flexible and shows a diminished capacity to dynamically reconfigure over time. In other words, although connectivity strength may remain within a nominal range, its ability to support diverse and adaptive network states is compromised. This constrained dynamical behavior implies that the network tends to settle into fewer, more stereotyped configurations, reflecting a reduced tendency toward dynamic transitions. Such a pattern may represent a distinct and previously underemphasized dimension of dysconnectivity in BD, wherein normal connectivity no longer facilitates rich dynamical variability but instead supports a more rigid and less adaptive functional network organization.

Similarly, H, derived from the eigenvalue spectrum of the adjacency matrix, indexes the stability and vigor of synchronization across the network[37,54,56,132]. The observed reduction of H in the theta band suggests that functional coupling in BD is less dynamically stable. Previous work has shown that energy-based measures are particularly sensitive to alterations in synchronization stability in neuropsychiatric and developmental conditions[37,54,56,132]. Within this framework, the present findings support the interpretation of BD as a condition characterized by unstable large-scale coordination, rather than a simple loss or gain of connectivity.

The ML results further reinforce this interpretation. Despite including both topological and dynamical features, the SVM classifier relied predominantly on entropy and energy to achieve optimal performance. Measures such as CD, CC, and Ef contributed relatively little to classification accuracy. This pattern indicates that how connections fluctuate and organize dynamically carries more diagnostic information than how they are arranged topologically. Such findings are consistent with recent arguments in network neuroscience that static connectome descriptions may overlook critical disease-related information embedded in temporal and spectral dynamics[103,107,133].

### Regional hub alterations and involvement of large-scale brain networks

Beyond whole-FBN alterations, our regional centrality findings provide important insight into which meso-scale FBNs are preferentially affected in BD and how network influence is redistributed across cortical systems. Specifically, we observed enhanced EigC in the right dorsolateral prefrontal cortex (BA 8) in BD, accompanied by reduced centrality in a set of temporal and limbic regions including BA 20, 21, 22, 28, and 36. These regions collectively span key components of the central executive network (CEN), default mode network (DMN), and limbic–temporal memory systems, suggesting a disrupted balance between cognitive control and affective–mnemonic processing in BD.

The increased centrality of BA 8, a core node of the dorsolateral prefrontal cortex (DLPFC), points to an enhanced influence of executive control regions within the FBN. The DLPFC is critically involved in cognitive control, attentional regulation, and top-down modulation of emotional responses, and has been repeatedly implicated in bipolar disorder across mood states[22,23,134–136]. Elevated hubness in this region may reflect a compensatory mechanism during the euthymic phase, whereby frontal control systems exert greater influence to maintain mood stability and cognitive regulation. However, excessive reliance on frontal hubs may also indicate an imbalanced network configuration, in which control-related regions dominate information flow at the expense of more distributed and flexible processing.

Conversely, the reduced centrality observed in temporal and limbic regions, including inferior and middle temporal gyri (BA 20, 21), Wernicke’s area (BA 22), entorhinal cortex (BA 28), and perirhinal cortex (BA 36), suggests diminished integration of regions that play central roles in emotional processing, semantic memory, and affective context representation. These areas are closely linked to the DMN and medial temporal memory system, which have been shown to exhibit abnormal connectivity and dysregulated interactions with frontal networks in BD[2,27,137–140] .Reduced EigC in these regions implies that they are less embedded within influential network neighborhoods, potentially limiting their contribution to large-scale information integration.

Once again, these regional effects were most pronounced in theta-band networks derived from nonlinear connectivity measures, underscoring the role of frequency-specific and nonlinear dynamics in shaping hub organization. Theta oscillations are known to support long-range coordination between prefrontal and limbic systems and are essential for cognitive–affective integration[104,141,142]. Altered theta-band hub structure therefore aligns with the broader interpretation that BD involves impaired coordination between executive and emotional systems, even during euthymic states.

## Conclusion

This study demonstrates that functional brain network dysconnectivity in euthymic BD is best characterized by alterations in network dynamics and complexity, rather than by global static topological reorganization. Individuals with BD showed reduced CD, lower S, and decreased H, primarily in theta and alpha1 bands, indicating diminished synchronization stability and a constrained repertoire of functional network states. After controlling density using null models, classical topological measures no longer differentiated groups, suggesting that global topology remains relatively preserved during euthymia.

At the regional level, altered hub organization revealed increased influence of frontal executive regions and reduced centrality in temporal–limbic areas, implicating disrupted interactions among meso-scale control and affective–mnemonic networks. Importantly, ML analyses confirmed that dynamic and information-theoretic features, particularly S and H, provided the strongest discrimination between groups. Together, these findings support a view of BD as a condition marked by impaired large-scale neural dynamics and reduced network flexibility, highlighting the potential of dynamic EEG-based graph measures as biomarkers.

### Limitations

The relatively small sample size may limit generalizability, although permutation testing and cross-validated ML were used to mitigate this issue. All patients were assessed during the euthymic phase, preventing conclusions about state-dependent network changes across mood episodes. Additionally, only undirected connectivity measures were examined, precluding inferences about directional information flow. Future studies using larger samples, higher-density recordings, longitudinal designs, and directed or time-resolved connectivity analyses are needed to extend these findings.

## Acknowledgments

The authors thank all participants for their time and cooperation in this study. This research received no specific grant from funding agencies in the public, commercial, or not-for-profit sectors.

